# Hematophagous triatomine bugs feed also on plants and express functional amylase

**DOI:** 10.1101/2023.03.03.530934

**Authors:** Jean-Luc Da Lage, Alice Fontenelle, Jonathan Filée, Marie Merle, Jean-Michel Béranger, Carlos Eduardo Almeida, Elaine Folly Ramos, Myriam Harry

## Abstract

**BACKGROUND:** Blood feeding is a secondary adaptation in hematophagous bugs that ancestrally feed on plants. Many vector proteins are devoted to cope with the host’s defenses and to process the blood meal. In contrast, one can expect that some digestive enzymes devoted to phytophagous diet were lost during or after this peculiar adaptation. And yet, in many strictly hematophagous arthropods, alpha-amylases genes, coding the enzymes that digest starch from plants, are still present and transcribed, including in the blood-sucking bug *Rhodnius prolixus* and its related species *R. robustus* (Hemiptera, Reduviidae, Triatominae). Triatominae bugs are vectors of *Trypanosoma cruzi*, the causal agent of Chagas’disease. Besides the parasitic human infection by the vector-borne route via an exposition to infected feces, an oral route is documented by the ingestion of contaminated food or juices made from palm fruit trees.

**METHODOLOGY/PRINCIPAL FINDINGS:** We hypothesized that retaining alpha-amylase could be advantageous if the bugs happen to consume occasionally plant tissues. To this end, we surveyed hundreds of gut DNA extracts from the sylvatic species *R. robustus* caught on palm trees to detect traces of plant meals. We found plant DNA in over 8 % of the *R. robustus* samples, mostly the palm tree *Attalea speciosa*. Moreover, we showed that the *R. robustus* alpha-amylase retained normal amylolytic activity.

**CONCLUSIONS:** Preserving alpha-amylase function could be an important way of optimally harness plant substrates, and plant feeding could be a way for bridging the gap between two blood meals. Our data indicate that plants are a common and yet underestimated food source in the wild for Triatomine.

**Author Summary:** Adaptation to a specific diet is often accompanied by metabolic, behavioral, physiological changes and hence by genetic changes like gene family expansion, gene losses or gains. In blood-sucking insects some adaptive features such as salivary components acting against blood clotting are known. However, it is intriguing that a digestive enzyme, alpha-amylase, which digests starch, is conserved in those animals, because blood does not contain starch nor related glucose polymers. This is the case in the blood-sucking bugs of the *Rhodnius* genus (Hemiptera, Reduviidae), which are vectors of the Chagas’disease, an important health issue in Latin America. In this study, we evidence for the first time that sylvatic bugs *R. robustus* also consume plant tissues in the wild. We detected by PCR performed on DNA from digestive tract that a significant number of wild-caught individuals harbored plant DNA, especially from *Attalea* palm trees, on which they used to nest. We showed that the amylase enzyme is normally active on starch. We suggest plant feeding could be a way for bridging the gap between two blood meals but might not be linked to nutritional distress.

## Introduction

Triatomine bugs (Hemiptera, Reduviidae), widespread in Latin America, transmit the parasite *Trypanosoma cruzi* (Kinetoplastidae, Trypanosomatidae), the causal agent of the Chagas’disease that affects 6-7 million people (WHO, 2016). Anthropogenic changes in natural ecosystems promote the home colonization by the sylvatic Chagas’disease vectors and increase the risk of human infection. In natural ecotopes triatomes feed on a large panel of vertebrates but some species exhibit niche specialization. Indeed, the genus *Panstrongylus* is predominantly associated with burrows and tree cavities, the genus *Triatoma* with terrestrial rocky habitats or rodent burrows while most of the *Rhodnius* species are associated with palms that could be used as an ecological indicator of epidemiological risk for Chagas’ disease (1). Among the genus *Rhodnius, R. prolixus* is of epidemiological importance for Chagas disease because this species has a wide distribution in central and northeastern Latin America and is largely a domesticated species. Thanks to chemical control campaigns, the prevalence of the disease due to this vector species has been considerably reduced (2). *Rhodnius robustus* is closely related to *R. prolixus* (3), (4) and widely distributed in the Amazon region. This species only sporadically visits human dwellings attracted by artificial light sources (5) which favors the transmission of *T. cruzi* to humans in particular because this species has a high parasitic prevalence (6).

Since Wigglesworth’ study (7), the triatomine bugs are considered to be strictly hematophagous, from the hatched nymphs to the mature male and female adults, as molting in nymph stages and oogenesis in adults are triggered by a blood meal. But, for the triatomine *Eratyrus mucronatus* living in large hollow trees, adults feed on porcupines (*Coendou prehensilis*) while it was reported that young instars can feed on the haemolyph of large arachnids (Amblypygi) which inhabit hollow trees (Gaunt & Miles, 2000). Among blood-feeding insects, the tsetse flies *Glossina spp*. (Diptera, Glossinidae) and the human louse *Pediculus humanus* (Phthiraptera, Anoplura) are considered strictly hematophagous. For others blood-feeding insects, only adults are hematophagous namely fleas (Siphonaptera), their larvae feeding on adult faeces (8), and flies, such as *Haematobia irritans* or *Stomoxys calcitrans* (Diptera, Muscidae), their larvae thriving in cattle dung (9, 10) but adults of *S. calcitrans* may collect sugar in the wild (9). Sometimes, only females are hematophagous such as mosquitoes (Diptera, Culicidae) and horseflies (Diptera, Tabanidae) whose females need a blood meal for egg maturation but have a mixed diet including nectar or plant consumption (11).

The salivary gland proteins injected by hematophagous insects when biting their prey include a cocktail of molecules necessary to cope with the blood composition and the prey’s immune defenses (12, 13). This involves inflammation modulators such as lipocalins, a class of which, nitrophorins, bind histamine. Because the prey’s immunity components might damage the predator’s organs, saliva also contains immunity modulators, such as complement inhibitors. Proteins with antihemostatic activity, inhibiting platelet aggregation or clotting, and vasodilators and proteases are also crucial. The heme from hemoglobin contains reactive iron ions that promote reactive oxygen species (ROS) and may alter cell membranes. The blood-feeding insects need therefore to detoxify blood molecules (14, 15).

Surprisingly, whereas blood is mainly rich in proteins and devoid of polysaccharides or oligosaccharides related to starch (maltooligosaccharides), a number of insect species considered strictly hematophagous do harbor alpha-amylase genes which are transcribed in salivary glands and in the digestive tract, as shown by numerous genome and transcriptome studies e.g. (16, 17, 18). This is the case in *Rhodnius prolixus* (19) and the related Triatomine species, the genomes or transcriptomes of which have been sequenced. We therefore wondered if this enzyme, that breaks down starch and maltooligosaccharides into smaller sugars, such as maltose, maltotriose and maltodextrins, has been recruited for another function in blood-sucking arthropods, or if it retained its original function. Interestingly, Diaz-Albiter et al (20) have shown that in a no-choice experiment in the laboratory, *R. prolixus* individuals were able to feed on tomatoes, although they were not able to achieve the life cycle. It is then legitimate to wonder if those insects can feed on plants in the wild. In this respect, it is noteworthy that, besides the parasitic human infection by the vector-borne route via an exposition to infected feces, an oral route is documented by the ingestion of contaminated food or beverages with infected triatomines or their feces (21, 22). The most *T. cruzi* oral cases were reported from Amazon region due to homemade juices from palm fruits contaminated by infected sylvatic insects, including the vector species *R. robustus* (23-25). Two explanation could be put forward, either the infected insects were passively transported with the fruit they were on, or the infected insects were actively attracted to the beverages being prepared and fell into. In both cases, a nutritional link between insects and fruits should be considered.

To this end, in the present study, using plant PCR detection, we have investigated the occurrence of plant feeding by sylvatic *Rhodnius robustus* individuals from Brazilian Amazon region sampled on palm tree species. We have also surveyed the enzymology of *Rhodnius* alpha-amylase to get an insight of its function.

## Methods

### Sampling

DNAs from *R. robustus* of various instars had been extracted previously (26) and kept frozen in the laboratory. The *R. robustus* individuals had been collected in 2008 and 2009 in Brazilian Amazonia in the Tapajós River region (Pará, Brazil) in three human communities (Saõ Tomé, Nova Estrela, Araipa Lago) on three palm trees species, *Attalea speciosa* (Babassu), *A. maripa* (Inaja) and *A. phalerata* (Urucuri). Gut DNAs were available for all instars except the smaller L1, L2 for which whole DNA has been extracted. Host-feeding sources have previously been identified by (27) using highly conserved primers for vertebrate *cytochrome b* gene (28). In this study, we first checked the quality of the DNA samples by amplifying a *cytochrome b* fragment with insect-specific primers (RhodCytbF and RhodCytbR) (29), suppl. Table S1). Negative-PCR samples were discarded.

The *Cimex lectularius* and *Rhodnius prolixus* strains were reared at the VITROME laboratory (IHU-Méditerranée Infection, Aix Marseille Université, Marseille, France).

### Detection of plant DNA

We then used 285 *R. robustus* individuals in order to detect DNA of plant tissues ingested by the insects. For detection of plants, we performed PCR on nuclear *18S* RNA gene fragments. Our attempts with classical chloroplast markers (30) or with internal transcribed spacers (ITS) (31) were not successful. Since *18S* is relatively well conserved among eukaryotes, we designed plant-specific primers (suppl. Table S1). PCR was carried out using the GoTaq polymerase (Promega). Amplification conditions are indicated in suppl. Table S1. Maize and apple DNAs were used as positive controls. Successfully amplified DNA fragments were sequenced in an ABI 3130 Sanger sequencer and the plant source was searched for by BLASTN (32) against the GenBank nr library. In 10 cases, where several sequences were mixed (sometimes including contaminants), the *18S* PCR products were purified (Nucleospin, Macherey Nagel) and cloned into the pCR2-Dual promoter plasmid (Invitrogen). Several clones were then sequenced to identify the different sequences in the PCR mixture.

### Amylase genes in hematophagous arthropods

We investigated the presence of alpha-amylase genes in insects and ticks considered as strictly hematophagous using TBLASTN search (32) in the NCBI nucleotide databases (genomes, nr, TSA, SRA). The query sequences are given in suppl. Table S2. The *R. robustus* alpha-amylase sequence was retrieved from the draft genome of this species (M. Merle, unpublished data).

### Phylogeny of amylase genes in Hemiptera

In order to get an overview of the presence of alpha-amylase genes in Hemiptera, we used the protein sequence of *R. robustus* as a TBLASTN query on hemipteran genomes and transcriptomes available at the NCBI (December, 2022). Sequences with acceptable length (66 species retrieved) were aligned with MUSCLE (33) (502 amino acid positions) and a maximum likelihood phylogenetic tree was computed at the iQ-TREE web server (34) with default parameters and 100 ultrafast bootstraps, and drawn using FigTree (http://tree.bio.ed.ac.uk/software/figtree/).

### Enzymology of R. robustus *alpha-amylase*

The alpha-amylase protein of *R. robustus* was produced in baculovirus-Sf9 cell system at the academic facility of the IGBMC (Illkirch, France). A dedicated peptide tag was added for the purification process. The pBacpak8 vector containing the amylase sequence in fusion with a Cterminal flag tag was synthetized by Genescript. 2 × 10^6^ of Sf9 (*Spodoptera frugiperda*) cells were co-transfected with 500 ng of *Bsu36I*-linearized BAC10:KO1629 DNA and 2 μg of pBacpak8-amylase with the calcium phosphate precipitate method and incubated at 27°C for 10 days. Harvested baculovirus were then tested, amplified and used to infect 3×0.5 l of Hi5 (*Trichoplusia ni*) cell culture for protein production. After 5 days of infection, the protein was purified from the medium using anti-flag M2 affinity gel (Merck). The protein was eluted from the beads with PBS + 10% glycerol + 0.5 mg/ml flag peptide. The protein was finally injected in a gel filtration column and eluted with PBS + 1mM TCEP. The sequence of the protein is given in suppl. Fig S1.

The amylolytic activity and parameters (*k*_cat_, K_m_, optimum pH, optimum temperature) were measured using the method of Bernfeld (35). The enzyme was diluted to 50 ng/μl (ca. 1 μM) and 20 μl of diluted enzyme was added to 200 μl of substrate solution and incubated at 37°C (except for the temperature curve). For the pH curve, soluble starch (Sigma 33615) was dissolved at 1% w/v in either acetate buffer (20 mM Na acetate, 20 mM NaCl, 1 mM CaCl_2_) for pH 3 to 5.5, MES buffer (20 mM MES, 20 mM NaCl, 1 mM CaCl_2_) for pH 6 and 6.5, HEPES buffer (20 mM HEPES, 20 mM NaCl, 1 mM CaCl_2_) for pH 7 and 7.5, Tris buffer (20 mM TRIS, 20 mM NaCl, 1 mM CaCl_2_) for pH 8 to 9.5. Finally, for *k*_cat_ estimate and temperature curve, acetate buffer at pH 4.5 was used at 1% w/v starch, and at various concentrations for the K_m_ curve, with 5 mn incubation. The K_m_ value was estimated from a non-linear regression using the Michaelis-Menten equation. The *k*_cat_ for glycogen (1%) was also estimated. We also assayed possible alpha 1,4 glucosidase activity using *p*-nitrophenyl-maltoside (Sigma N5885) as a substrate at 5 mg/ml in acetate buffer pH 4.5, with 2 μl of enzyme at ca. 1.5 μM in 200 μl of substrate. The release of *p*-nitrophenol at 37°C was recorded every 5 mn for one hour by the increasing absorbance at 405 nm. Measures were made in duplicates or triplicates.

Alpha-amylase activity was also recorded from laboratory-bred insects. Whole blood-feeding bed bug *Cimex lectularius* or head + thorax from *Rhodnius prolixus* were ground in 50 μl ice-cold water, then centrifuged 5 mn; 10 μl were spotted onto starch (0.5%)-agar plates containing 20 mM NaCl and 1 mM CaCl_2_, and incubated one hour at 37°C. Activity was revealed with Lugol as white spots on a blue background.

We tested the possible action of the alpha-amylase of *R. robustus* on various oligo-and polysaccharides and observed the products by thin layer chromatography, as described in (36). All substrates were dissolved at 5 mg/ml in acetate buffer, pH 4.5 and incubated for 24h at 37°C (2 μl of enzyme at ca. 1.5 μM in 200 μl of substrate). Commercial references of the chemicals are as follows : soluble starch : Sigma 33615 ; amylopectin : Sigma 10120 ; glycogen : ICN 10839 ; maltotriose : Sigma M8378 ; trehalose Sigma T0167 ; β-cyclodextrin : TCI C0777 ; γ-cyclodextrin : TCI C0869 ; dextran : Sigma 31390 ; nigeran : Sigma N8634 ; maltopentaose : Sigma 47873 ; mannan : Sigma M7504 ; maltohexaose 47876 ; saccharose : Prolabo 27480 ; raffinose : Euromedex UR1030 ; cellotetraose : Sigma C8286 ; heparin : Sigma H3393.

## Results

### Detection of plant DNA in R. robustus

We performed PCR amplification on gut or whole-body DNAs in order to detect plant material. As we faced likely contaminations by plant material from our environment, we decided to discard positive results when they were identified (with 98-100% sequence identity) as European plants possibly present around the laboratory (*Acer, Cedrus, Pinus, Quercus, Betula, Humulus, Fagus, Urtica*…) and to keep only tropical plants from Brazil. After correction for contaminated or dubious samples and using a high confidence identity (≥ 99.5 %) on at least 500 bp, we obtained positive results for 24 *R. robustus* samples over 285 (8.4 %, Table 1). Eighty-seven percent (21/24) of the samples exhibited *18S* amplicons that matched with the palm tree *Attalea speciosa* (Arecaceae) identification. Three samples (22, 122, 915) harbored two plant species. Among the samples with plant DNA, two-thirds (16/24) were not morphologically supplied with blood when they were sampled. Therefore, there does not appear to be a relationship between insect blood nutritional status and the presence of plant DNA.

**Table 1:** Identification of plants ingested by *R. robustus* using *18S* primers. G: gut; WO: whole organism, B: presence of fresh blood in the digestive tract, NB: no fresh blood in the digestive tract.

| Sample number | Insect stage | Locality | Sampling | Nutritional status | Tissue used | plant identified | Identity / sequence length |
| --- | --- | --- | --- | --- | --- | --- | --- |
| 22 | M | Nova Estrela | *A. phalerata* | NB | G | *Attalea speciosa* | 99.6% / 766 bp |
|  |  |  |  |  |  | *Cecropia pachystachya* | 99.6% / 766 bp |
| 116 | L5 | Nova Estrela | *A. phalerata* | NB | G | *Attalea speciosa* | 99.6% / 551 bp |
| 122 | L6 | Nova Estrela | *A. phalerata* | NB | G | *Adenocalymma validum* | 99.9% / 817 bp |
|  |  |  |  |  |  | *Cecropia pachystachya* | 99.8% / 679 bp |
| 145 | L5 | Araipá Lago | *A. maripa* | NB | G | *Attalea speciosa* | 100% / 743 bp |
| 172 | L5 | São Tomé | *A. speciosa* | NB | G | *Attalea speciosa* | 99.9% / 727 bp |
| 173 | L5 | São Tomé | *A. speciosa* | NB | G | *Attalea speciosa* | 100% / 805 bp |
| 209 | F | São Tomé | *A. phalerata* | NB | G | *Attalea speciosa* | 99.8% / 616 bp |
| 211 | L5 | São Tomé | *A. phalerata* | NB | G | *Attalea speciosa* | 99.6% / 789 bp |
| 214 | L5 | São Tomé | *A. phalerata* | B | G | *Attalea speciosa* | 99.6% / 779 bp |
| 224 | L4 | São Tomé | *A. phalerata* | B | G | *Cecropia ficifolia* | 99.9% / 747 bp |
| 225 | L4 | São Tomé | *A. phalerata* | B | G | *Attalea speciosa* | 99.6% / 552 bp |
| 226 | L4 | São Tomé | *A. phalerata* | NB | G | *Attalea speciosa* | 99.9% / 783 bp |
| 248 | L3 | São Tomé | *A. phalerata* | B | G | *Attalea speciosa* | 99.8% / 662 bp |
| 249 | L2 | São Tomé | *A. phalerata* | B | WO | *Attalea speciosa* | 99.4% / 637 bp |
| 599 | L3 | São Tomé | *A. maripa* | NB | G | *Attalea speciosa* | 99.6% / 772 bp |
| 806 | L3 | São Tomé | *A. phalerata* | B | G | *Attalea speciosa* | 99.7% / 405 bp |
| 821 | L4 | Nova Estrela | *A. maripa* | B | G | *Attalea speciosa* | 100% / 709 bp |
| 850 | L5 | Nova Estrela | *A. phalerata* | B | G | *Cecropia pachystachya* | 99.5% / 796 bp |
| 857 | L2 | Nova Estrela | *A. phalerata* | NB | WO | *Attalea speciosa* | 100% / 818 bp |
| 869 | L5 | Nova Estrela | *A. phalerata* | NB | G | *Attalea speciosa* | 99.8% / 525 bp |
| 875 | L3 | Nova Estrela | *A. phalerata* | NB | G | *Attalea speciosa* | 100% / 534 bp |
| 877 | L3 | Nova Estrela | *A. phalerata* | NB | G | *Attalea speciosa* | 99.7% / 596 bp |
| 878 | L3 | Nova Estrela | *A. phalerata* | NB | G | *Attalea speciosa* | 99.9% / 828 bp |
| 915 | L2 | Nova Estrela | *A. speciosa* | NB | WO | *Attalea speciosa* | 99.6% / 566 bp |
|  |  |  |  |  |  | *Nephrolepis biserrata* | 99.7% / 785 bp |

### Amylase gene in hematophagous arthropods

We investigated the presence of amylase genes in transcripts and/or genes of strictly hematophagous insect and tick species (Table 2). For all blood-sucking bugs (Hemiptera, Reduviidae, Triatominae), tsetse flies (Diptera, Glossinidae) and ticks (Ixodidae, Argasidae), for which nuclear data are available, at least one alpha-amylase gene/transcript was found, including *Rhodnius, Triatoma* and *Panstrongylus* genus for the Triatominae, three *Glossina* species for tsetse flies, and *Amblyomma, Hyalomma, Rhipicephalus, Haemaphysalis, Dermacentor* and *Ornithodoros* genus for ticks. However, for the human louse *Pediculus humanus* and other blood-sucking lice (Phthiraptera, Anoplura) no sequence was retrieved.

**Table 2:**
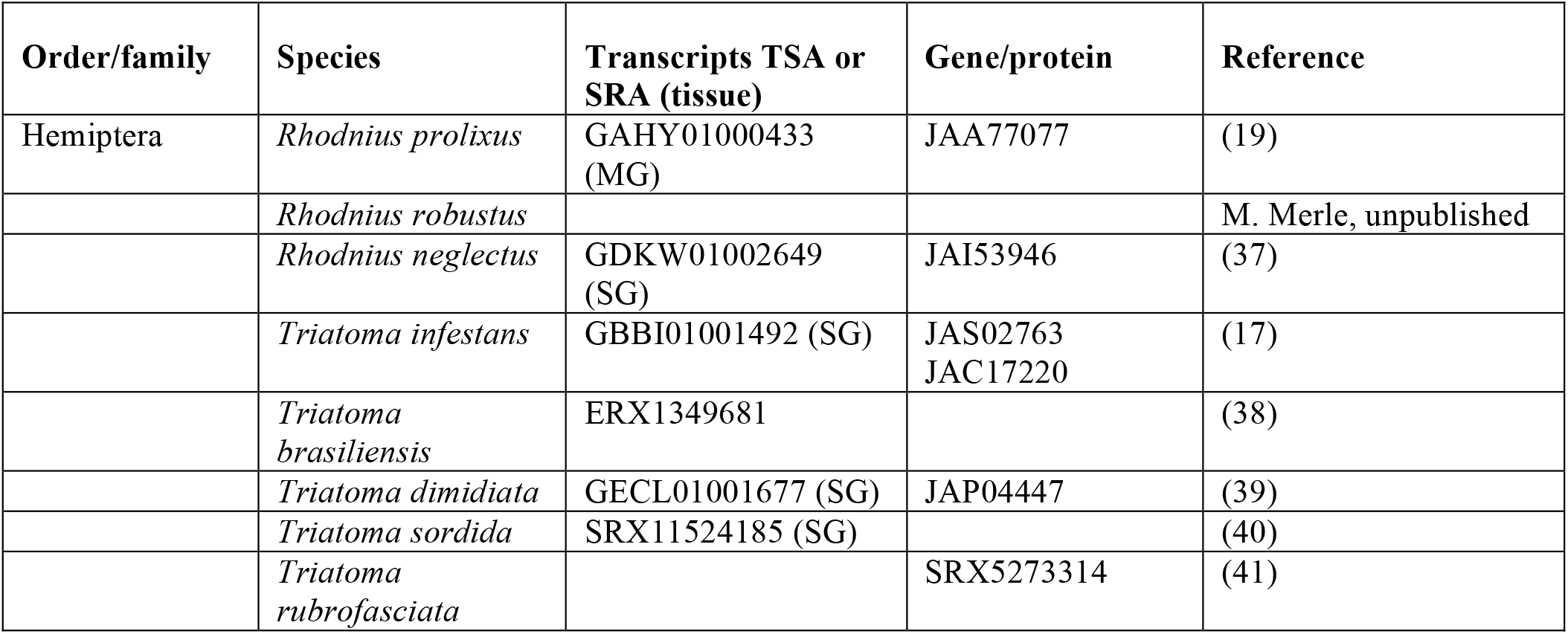
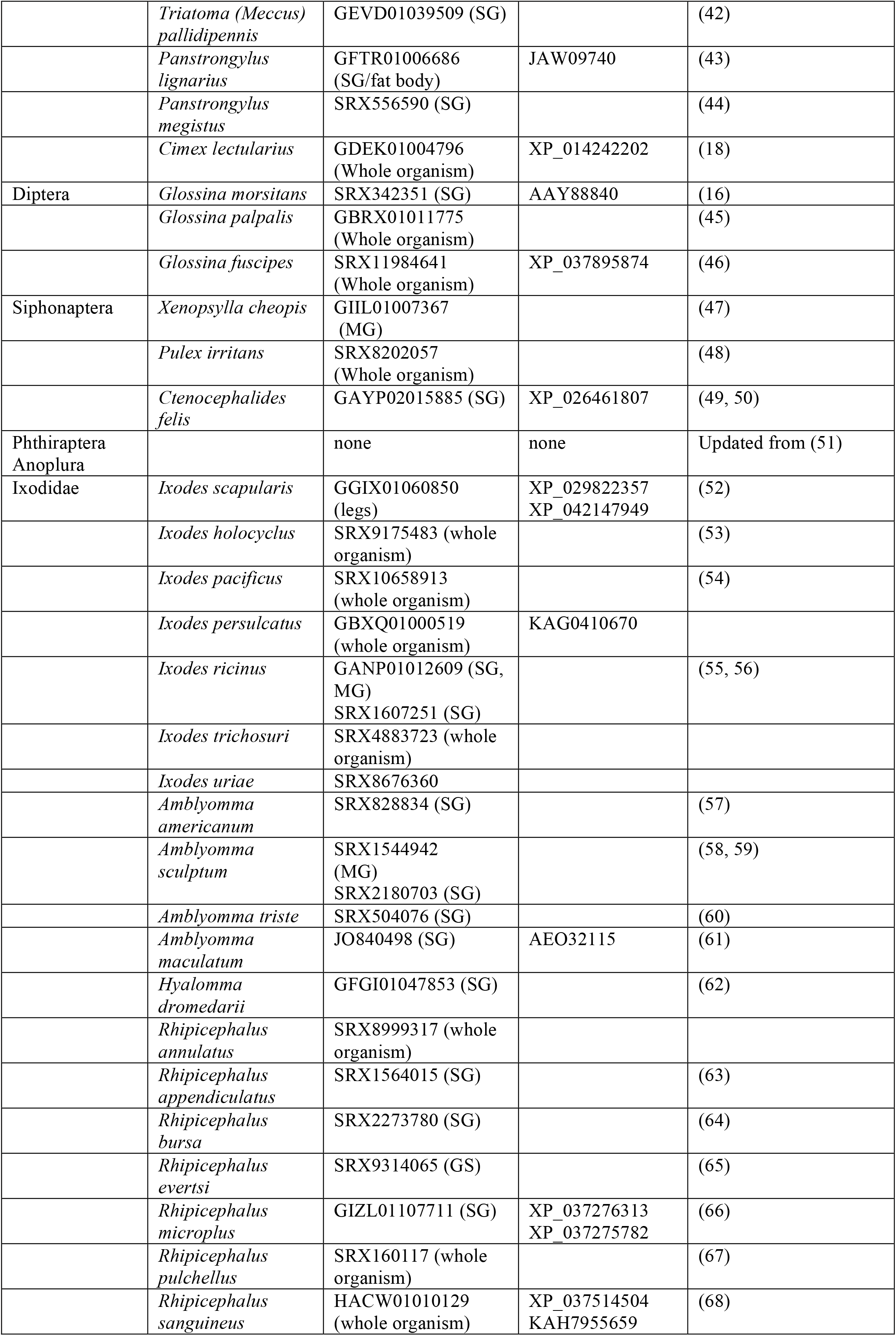
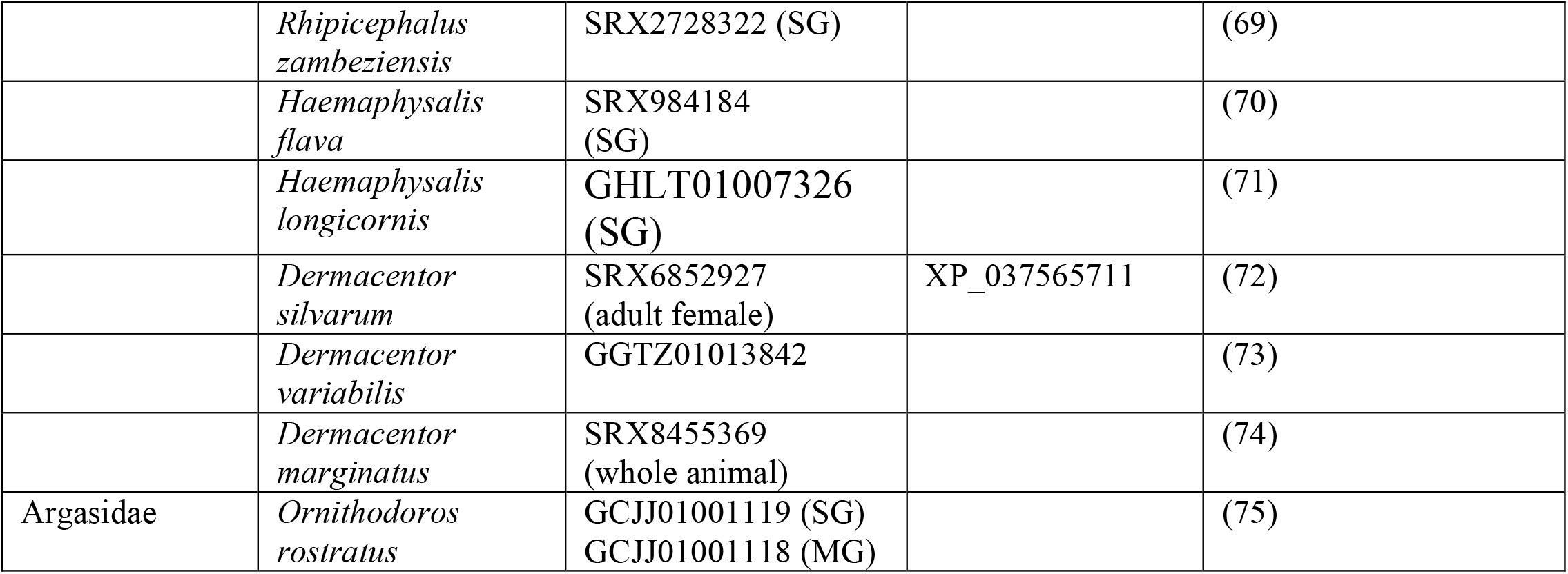
Occurrence of alpha-amylase transcripts and/or genes in strictly hematophagous species. SG : sialome (salivary gland) ; MG : midgut. TSA : Transcriptome Shotgun Assembly ; SRA : sequence read archive. The accession numbers are from GenBank. SRA and TSA databases were searched using TBLASTN with the *R. neglectus* protein sequence JAI53946 as a query for Hemiptera, *G. morsitans* AAY88840 for Diptera, Siphonaptera, and Phthiraptera, *I. scapularis* XP_029822357 for Ixodidae and Argasidae. The Phthiraptera Anoplura searched were *Pediculus humanus* (genome) and SRA data from *Polyplax serrata* (SRX8978347), *Hoplopleura arboricola* (SRX8976670), *Neohaematopinus pacificus* (SRX8976664 and SRX8976650), *Echiniphthirus horridus* (SRX4966554 and SRX4966544), *Phthirus pubis* (SRX2405464), *Phthirus gorillae* (SRX2405463), *Antarctophthirus microchir* (SRX2405454).

### Amylase gene phylogeny in Hemiptera

Our search in the whole Hemiptera (suppl. Table S3 and Fig 2) shows that alpha-amylase is widely conserved in hemipterans including Triatominae and Cimicidae whereas it was lost in particular parasitic sap feeders such as aphids and mealybugs. Note that it is likely that in a number of species, several genes exist. Two amylase sequences of *Halyomorpha halys* were included as an example.

**Figure 1:**
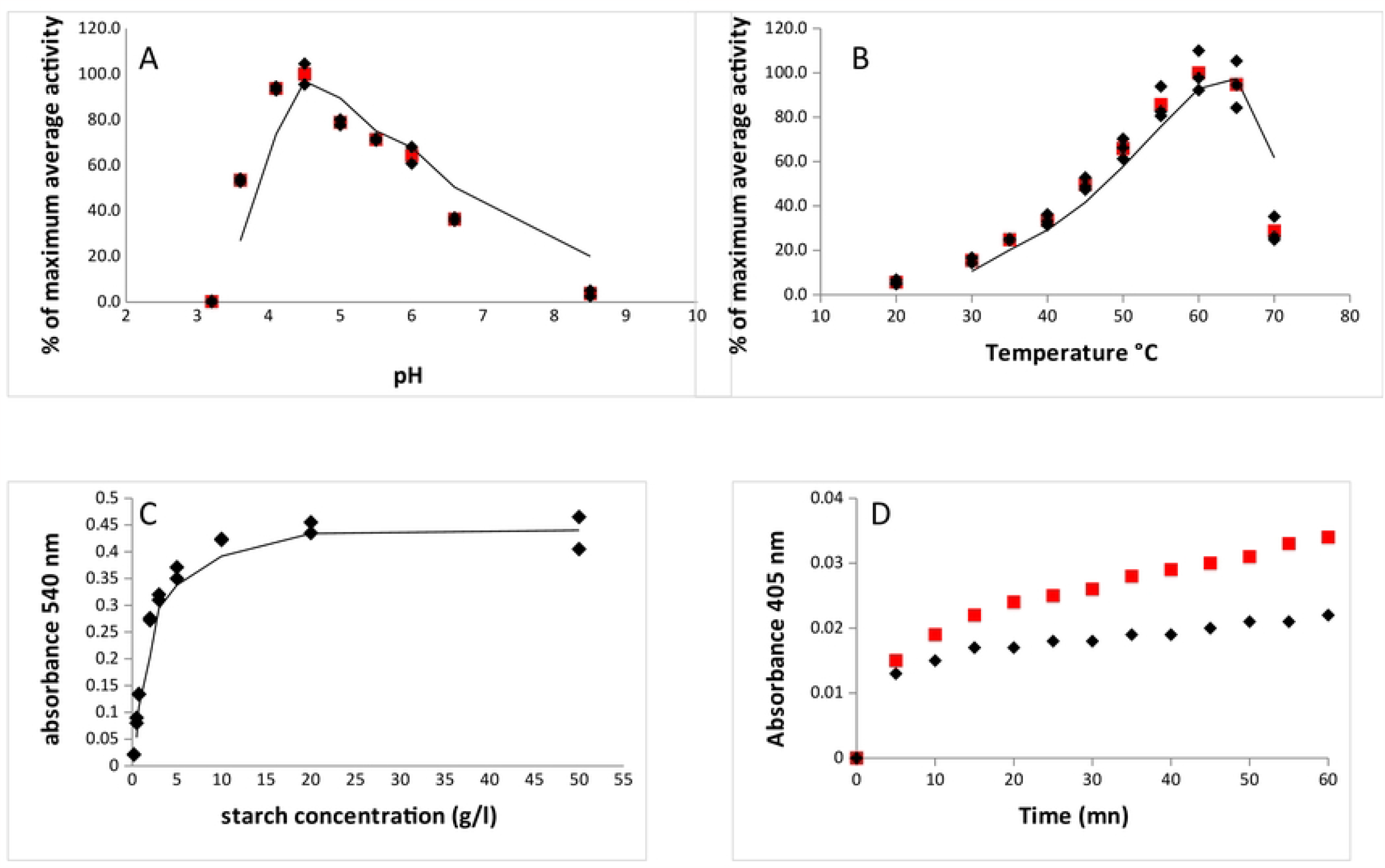
Enzymological characterization of the purified alpha-amylase of *R. robustus*. A) alpha-amylase activity as a function of pH on 1% starch at 37°C ; B) alpha-amylase activity as a function of temperature at the optimum pH 4.5 on 1% starch, 5 mn incubation; C) alpha-amylase activity as a function of starch concentration at 37°C and pH 4.5 for K_m_ estimate. Black diamonds are the individual replicates, red squares are averages of 2-3 replicates. D) measure of alpha-1,4 glucosidase activity on *p*-nitrophenylmaltoside. Black diamonds: incubation without enzyme (autolysis) ; red squares : incubation with enzyme.

**Figure 2:**
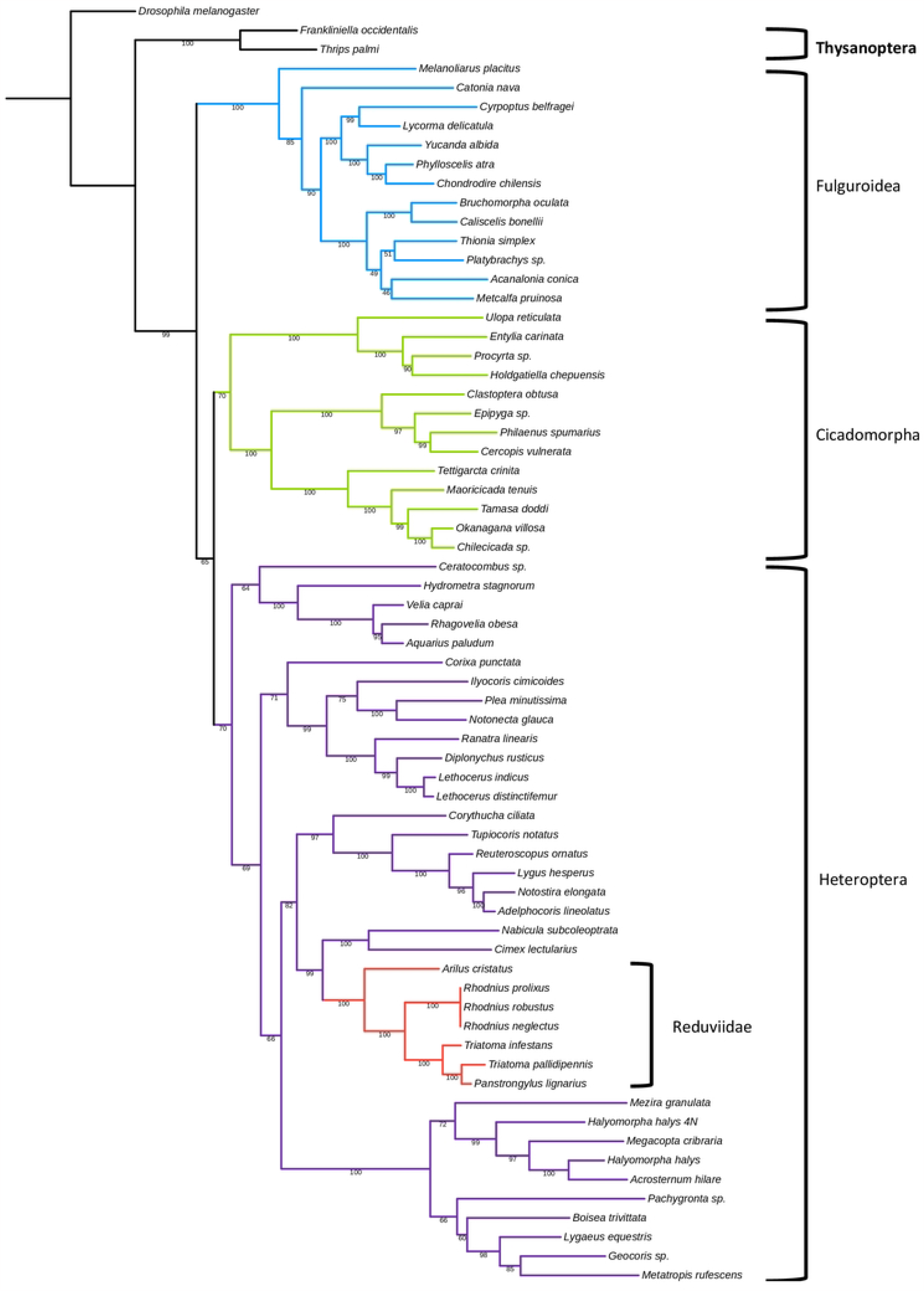
Maximum likelihood phylogenetic tree of hemipteran alpha-amylase protein sequences, computed by the iQtree web server (34) with default parameters. The tree was rooted with *D. melanogaster*. Purple : Heteroptera, with Reduviidae in red ; green : Cicadomorpha ; blue : Fulgoroidea. Extra groups are in black.

### Enzymological characteristics of the R. robustus *amylase*

The mature alpha-amylase of *R. robustus* was produced *in vitro* and purified from cultured insect cells and is considered to be similar to the natural enzyme, although there is a C-terminal tag used for purification purpose. We carried out classical enzymological measures. The pH, temperature and K_m_ curves are shown on Fig 1A, 1B, 1C, respectively. The optimal pH observed is 4.5 and the optimal temperature 60°C. The catalytic activity *k*_cat_ at 37°C was estimated to be 81 s^-1^ on starch and 58.5 s^-1^ on glycogen, which is an equivalent of starch in animal tissues. This is a moderate value, compared to the well-studied *D. melanogaster* amylase, whose *k*_cat_ is ca. 300-500 s^-1^ at the lower temperature 25°C (76). The K_m_ is 1.71g l^-1^ on starch. The catalytic efficiency *k*_cat_/K_m_ is therefore 47.4 s^-1^ g^-1^ l. Thus, we show that the *R. robustus* alpha-amylase (and *R. prolixus*, which has the same sequence) is active and does exhibit its original function of hydrolyzing starch. In addition, it shows a slight alpha-glucosidase activity, releasing low amounts of glucose from short-chain oligosaccharides such as maltotriose, or *p*-nitrophenol from its proxy *p*-nitrophenylmaltoside (Fig 1D and suppl. Fig S2). We also checked that there was an alpha-amylase activity *in vivo*. Supplementary Fig S3 shows that blood-feeding insects *R. prolixus* or *C. lectularius* bred in the laboratory, exhibit slight amylase activity, as evidenced by decoloration of starch-iodine complex after incubation with raw soluble extracts. We also surveyed the possible action of *R. robustus* alpha-amylase on various polysaccharides, with no evidence of breakdown of oligo- or polysaccharides other than maltooligosaccharides, which suggests that this enzyme has a classical amylolytic activity (suppl. Fig S2).

## Discussion

Blood feeding is not the ancestral state in bugs, but rather phytophagy, with possible intermediate states such as predation leading to hematophagy, which evolved independently in Reduviidae and Cimicidae (77). Accordingly, Hwang and Weirauch (78) showed that the generalist predatory feeding strategy is ancestral for Reduviidae, the assassin bugs. But, in the New World, members of the tribe Harpactorini are associated with some plants providing in addition to hosting the arthropod prey species the assassin bugs feed on, sugary or proteinaceous secretions produced in extrafloral nectaries or other structures (79, 80). In line with this, alpha-amylase has been detected in predaceous Reduviidae (reviewed in (81)). In this context, our primary goal in this study was to explain the persistence of a gene such as alpha-amylase, that breaks down starch and maltooligosaccharides into smaller sugars, in a strict blood-feeding insect derived from predatory ancestor. Our results show that alpha-amylase is widely conserved in hemipterans including hematophagous Triatominae and Cimicidae, and was lost in particular parasitic sap feeders such as aphids and mealybugs. The alpha-amylase phylogenetic tree roughly fits to the known hemipteran phylogeny (82). Indeed, since this enzyme processes polysaccharides related to starch or glycogen, it is expected to be useless for a hematophagous diet. A possible explanation is that those insects are not strictly hematophagous and, consequently, by uptaking vegetal materials which often contain starchy components, they would benefit from the production of a functional starch-degrading enzyme. It would suggest that those insects may need, at least occasionally, the amylolytic function when they encounter and eventually ingest starchy substrates. In this case, if injected into the plant, the enzyme would help digesting the plant tissue before absorption through the stylet.

We surveyed a large sample of sylvatic *R. robustus* individuals for plant DNA detection. We successfully identified neotropical plants in over 8 % of the *R. robustus* samples. It is likely that this percentage of plant detection is underestimated, given that the DNA samples were old and had been thawed-refrozen many times. It is unlikely that the plant DNA could be contaminating material present on the cuticle, since in most cases, the DNA was extracted from midgut, not from whole animals. The plant most commonly identified in our samples was *Attalea* palm tree (87 %). This could be expected, since most of the *Rhodnius* species, including *R. robustus*, dwell in palm trees. The only recovery of *A. speciosa* from the *18S* identification while the specimens of our study were sampled from the three species *A. speciosa* (Babassu), *A. maripa* (Inaja) and *A. phalerata* (Urucuri), is explained by the fact that only this species of *Attalea* is listed in *18S* database.

It is not clear which plant tissues were consumed. At least, it is likely that fruits of *Attalea* are attractive for *Rhodnius*, because massive human contaminations by Chagas disease’s vector may occur when people eat palm fruits or consume palm fruit juices, which thus prove to have been contaminated by the bugs (22).

Plant feeding could be a way for bridging the gap between two blood meals since finding a blood source in the wild may be tricky leading to a nutritional status of the bugs often poor (83). This could be particularly true for *Attalea sp*., which, in general, bloom and bear fruit throughout the year (84). *Attalea* fruit is a drupe with a starchy mesocarp (the Babassu flour contains about 60 % w/w starch) (85), and not only the fruits are starchy, but also the stems of palm trees (86). Hence, preserving alpha-amylase function could be an important way of optimally harness plant substrates. In this respect, we have confirmed in this study that alpha-amylase of *R. robustus* is a functional amylolytic enzyme. The enzymological study showed specific data as an optimal pH relatively low (4.5) compared to other data available for Hemiptera (51) and a high optimal temperature (60°C) compared to e.g. 50-55°C for purified alpha-amylase of *Drosophila melanogaster* (87). To note, the optimal temperature would be lower if we used a raw extract, because the presence of proteases would interfere and decrease the integrity and activity of the amylase (e.g. (88, 89)).

Our finding of plant consumption by hematophagous bugs is the first evidence to our knowledge of such a feeding behavior in the wild for hematophagous bugs. This is not quite surprising. Diaz-Albiter et al (2016) (20) have shown that forced feeding on plants in the laboratory (cherry tomatoes in their study) was possible, and that the bugs survived and improved their physiological status compared to starved controls, although not able to achieve their life cycle. In addition, they observed that a sweet food supply would stimulate subsequent blood feeding. These authors suggested that occasional plant feeding would explain the commonly observed association of triatomine bugs with tropical plants. In this respect, the fact that many arthropod species considered as purely hematophagous such as e.g. Ixodidae ticks or the tsetse fly do have a transcribed alpha-amylase gene (Table 2) would suggest that they may occasionally use plant resource too. Those species are free-living hematophagous insects. Interestingly, Phthiraptera Anoplura (blood-sucking lice) which are, in contrast, permanently stuck to their host and for which blood is thus always available have no longer an alpha-amylase gene. Notably, Phthiraptera Amblycera (chewing lice), which feed mainly on epiderma or feathers, may have retained an alpha-amylase gene (Suppl. Table S3).

Also, plant feeding might not be linked to nutritional distress. Consistent with this, one-third of *R. robustus* specimens with DNA plant in their gut were blood engorged when sampled. While the enzymological parameters measured in our study show a classical case, this does not preclude the possibility that the amylase protein might have also a non-catalytic function and be involved in processes linked to the blood diet. In line with this, Mury et al. (15) have shown that an alpha-glucosidase in *R. prolixus* is involved in hemozoin processing, i.e. detoxication of the reactive haeminic iron from hemoglobin, a nice case of exaptation. Plant feeding might supply micronutrients or chemical molecules involved in blood digestion. In *Rhodnius prolixus*, the gut symbiotic bacteria *Rhodococcus rhodnii* provides their hosts with B-group vitamins lacking from ingested blood and *Wolbachia* could ensure or complement, the nutritional mutualism provided by the gut symbionts (Filée et al., https://www.biorxiv.org/content/10.1101/2022.09.06.506778). We can thus hypothesize that plant food may also be the source of new probiotics or nutrients able to compensate for the transient absence of some nutritional symbionts. Feeding in Triatomine could even be a more complex process involving plant resources. An experimental study showed that oral treatment with babassu mesocarp has a significant anti-thrombotic effect in mice related to an increase in the ability of macrophages to produce nitric oxide (NO) and a slow coagulation process (90). These mechanisms are to be compared to those induced by different salivary proteins during blood-feeding. It could be hypothesized that plant molecules could be taken from the palm tree and reused by the vector during its food intake.

Additional studies are needed to assess the effect of ingesting plant material not only on the microbiome of the vector, including its symbionts, but also on the presence of the parasite *T. cruzi*, particularly in *R. robustus* which has a high parasitic prevalence.

This study constitutes an important advance in the questioning of the association of vectors of the genus *Rhodnius* with palm trees. Other studies are to be undertaken to search for the presence of plant material in other species of Triatominae.

### Ethical statement

This study strictly followed the ethical codes of Brazil. The DNA samples used in this study come from a previous sampling carried out in full compliance with the Brazilian laws (IBAMA authorization no.: 16485-2, 27/06/2008) during the field trip of the International Research Program, PLUPH project, supported by the Canadian Global Health Research Initiative and coordinated by M. Lucotte (Université du Québec, Montréal, Canada) and J. Drummond (Universidade de Brasilia, Brazil).

## Acknowledgments

We thank the platforms and common services of IGBMC (Illkirch, France), in particular the baculovirus facility, for amylase synthesis. We thank M. Quartier and F.B.S. Dias for the DNA extraction performed in 2010, M. Tenaillon and A. Cornille for supplying control DNA from maize and apple, respectively.

---

**Supplementary Figure S1:**
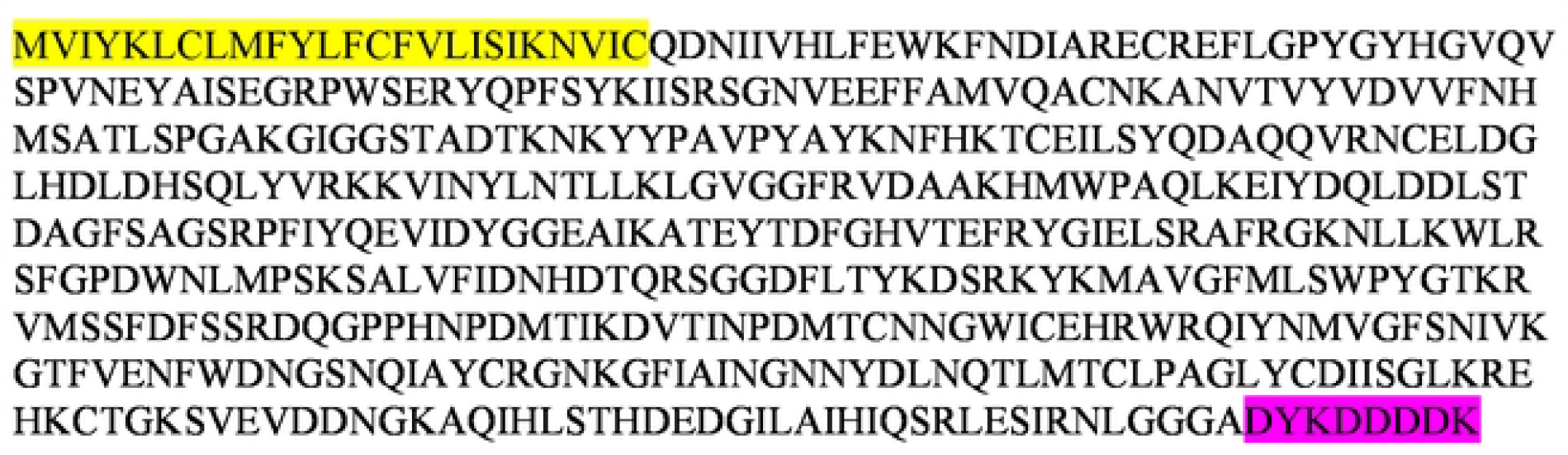
Sequence or the alpha-amylase protein of *Rhodnius prolixus*. Yellow highlight, native signal peptide ; pink highlight, purification tag.

**Supplementary Figure S2:**
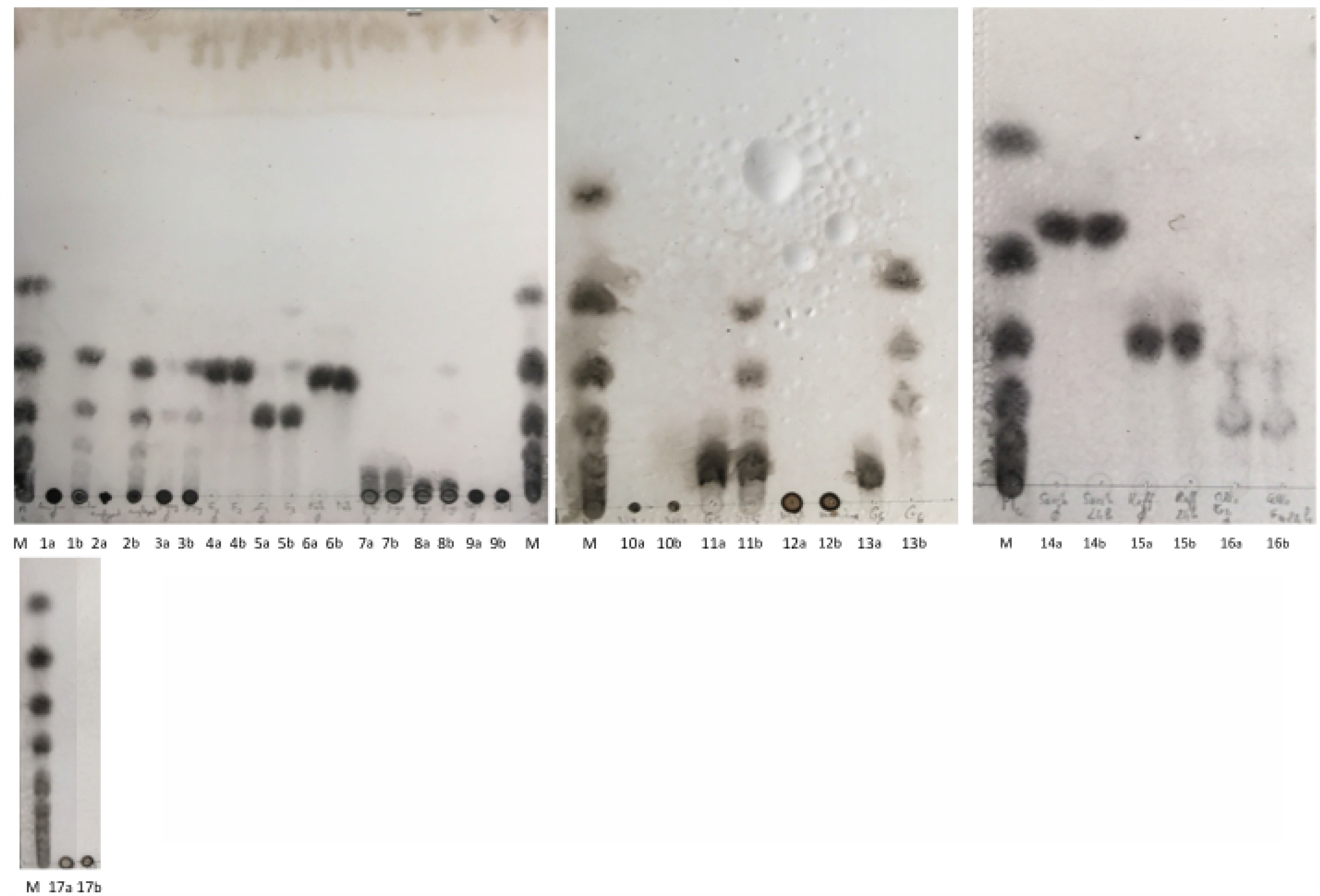
Thin layer chromatography of the action of *R. robustus* alpha-amylase on various sugars. For each spot, 5 × 3 μl were dried onto the silica plate and a negative control (a) incubated without enzyme was spotted next to the enzyme-treated sample (b). M : maltooligosaccharide ladders ; 1 : starch ; 2 :amylopectin ; 3 : glycogen ; 4 : maltose ; 5 : maltotriose ; 6 :trehalose ; 7 : β-cyclodextrin ; 8 : γ-cyclodextrin ; 9 : dextran ; 10 nigeran ; 11 : maltopentaose ; 12 : mannan ; 13 : maltohexaose ; 14 : saccharose ; 15 : raffinose ; 16 : cellotetraose ; 17 : heparin.

**Supplementary Figure S3:**
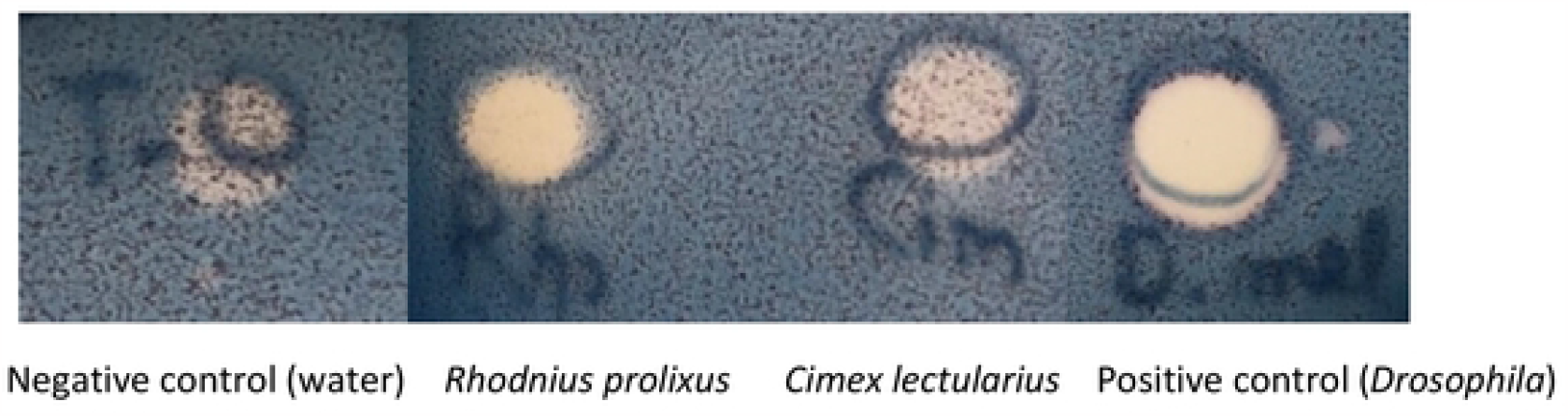
Starch-iodine assay of soluble extracts of *Rhodnius prolixus* (head and thorax) and *Cimex lectularius* (whole body). A larva of *Drosophila melanogaster* was used as a positive control.

